# VSV-based vaccine provides species-specific protection against Sudan virus challenge in macaques

**DOI:** 10.1101/2022.10.27.514045

**Authors:** Andrea Marzi, Paige Fletcher, Friederike Feldmann, Greg Saturday, Patrick W. Hanley, Heinz Feldmann

## Abstract

The ongoing Sudan virus (SUDV) outbreak in Uganda highlights the need for rapid response capabilities against emerging viruses with high public health impact. While such countermeasures have been established for Ebola virus (EBOV), they unfortunately do not exist for SUDV or any other human-pathogenic filovirus.

Here, we describe the generation and characterization of the vesicular stomatitis virus (VSV)-based vaccine VSV-SUDV and demonstrate the protective efficacy following a single-dose vaccination against lethal SUDV infection in nonhuman primates (NHPs). As we repurposed NHPs from a successful VSV-EBOV vaccine efficacy study, we further demonstrate that VSV-SUDV can be used effectively in individuals previously vaccinated against EBOV. While the NHPs developed cross-reactive humoral responses to SUDV after VSV-EBOV vaccination and EBOV challenge, cross-protection was limited emphasizing the need for the development of specific countermeasures for each human-pathogenic ebolavirus. Additionally, our data provides evidence that while previous VSV-EBOV immunity is boosted after VSV-SUDV vaccination, it has only limited impact on the immunogenicity and protective efficacy of VSV-SUDV vaccination important for frontline outbreak workers.

## Introduction

The ongoing Sudan virus (SUDV) outbreak in Uganda^1^ has focused our attention back on filoviruses, yet away from the well-known Ebola virus (EBOV). The current outbreak started with a 24-year-old male who was diagnosed on September 19 after visiting several health clinics^1^. Cases are currently reported from the Buyangabu, Kampala, Wasiko, Kagadi, Kyegegwa, Mubende, and Kassanda districts in Central and West Uganda approximately 160 kilometers west of Kampala^2^. There is on-site laboratory testing and medical support to identify and manage case patients. However, specific treatment and vaccines are currently not available to assist with outbreak management^1^.

Today six distinct species of ebolavirus have been described^3^ of which *Zaire*, *Sudan*, *Bundibugyo*, and *Taï Forest ebolaviruses* are known causes of human hemorrhagic disease^4^. SUDV, the single virus member in the *Sudan ebolavirus* species, was co-discovered with EBOV in 1976 during an outbreak of viral hemorrhagic disease (now designated Sudan virus diseases (SVD)) in what is now The Republic of South Sudan. SUDV remerged in The Republic of South Sudan in 1979 and 2004 causing smaller SVD outbreaks. In 2000/01, SUDV emerged in Gulu, Uganda causing the so far largest SVD outbreak on record with 425 cases and a case fatality rate of 53%^5^. This was followed by smaller outbreaks in Uganda in 2011, 2012, and 2012/13^6^. As of 25 October 2022, the ongoing SVD outbreak in Uganda accounts for 109 confirmed cases^7^. The overall case fatality rate of SUDV infections is ~50% and the current outbreak does not seem to differ significantly. Apart from outbreaks caused by SUDV, Uganda has previously reported outbreaks caused by Bundibugyo virus in 2007 and EBOV in 2019^6^ (Table S1) which show average case fatality rates of 25% and 50%^8^, respectively^9^.

There are currently two vaccines licensed for Ebola virus disease (EVD) caused by EBOV, the single shot VSV-EBOV vaccine (Ervebo®, Merck) and the Ad26.ZEBOV/MVA-BN-Filo prime-boost approach (Zabdeno^®^ and Mvabea^®^, Johnson & Johnson). While VSV-EBOV would be immediately available, it is unlikely to cross-protect against SUDV infections due to antigenic differences of the viral glycoprotein (GP)^10^. Ad26.ZEBOV/MVA-BN-Filo may be protective as the MVA-BN-Filo component includes a SUDV GP antigen, but clinical data is not available. As of today, multiple vaccine candidates targeting SUDV or several filoviruses including SUDV are in preclinical development^11^. The vaccines are mainly based on platforms that have previously been investigated for EBOV-specific vaccines. The platforms include viral vectors (i.e. human and chimpanzee adenoviruses, VSV, human parainfluenza virus type 3, rabies virus, modified vaccinia Ankara (MVA), Venezuelan equine encephalitis virus), virus-like particles, protein subunit and DNA as outlined in WHO’s Landscape of SUDV vaccine candidates^11^. Those approaches with promising efficacy in preclinical nonhuman primate (NHP) studies are listed in Table S2. Only three candidate vaccines have been or are currently in phase 1a/b clinical trials addressing toxicity and immunogenicity: the chimpanzee adenovirus serotype 3 (cAd3) expressing the SUDV GP (www.clinicaltrials.gov; NCT04041570 and NCT04723602), the chimpanzee adenovirus ChAdOx1-BiEBOV expressing the EBOV and SUDV GPs (www.clinicaltrials.gov; NCT0509750 and NCT05301504), and a DNA-based vaccine expressing Marburg virus GP, EBOV GP and SUDV GP^12^ which is no longer being pursued. Thus, together with the Ad26.ZEBOV/MVA-BN-Filo, there are three candidates to consider for potential use in the current SUDV outbreak.

We have developed a VSV-based SUDV-specific vaccine candidate expressing the SUDV-Gulu strain GP instead of the VSV glycoprotein (G) (here designated VSV-SUDV). SUDV-Gulu is the causative strain of the biggest recorded SVD outbreak from 2000/01 in Uganda (Table S1) and phylogenetically closer than the original SUDV-Boniface strain to the current outbreak strain^13^. The vaccine vector is based on the VSV G replacement approach and, thus, in principle identical to VSV-EBOV. Here we show in the cynomolgus macaque model that a single dose vaccination with VSV-SUDV is uniformly protective against SUDV challenge, that pre-existing EBOV immunity does not affect the protective efficacy of VSV-SUDV against SUDV challenge, and that pre-existing VSV-EBOV immunity does not protect against SUDV challenge despite cross-reactive immune responses. This vaccine candidate has successfully completed preclinical evaluation and is ready to be moved into clinical studies.

## Results

### Vector and study design

VSV-SUDV was generated according to previously published methods ^14,15^. The SUDV-Gulu GP open reading frame was inserted into the VSV backbone replacing the VSV G (Fig. S1A top). SUDV GP expression in VSV-SUDV-infected cells was confirmed by immunoblot (Fig. S1B). For *in vivo* efficacy testing of VSV-SUDV we used survivors of a previous VSV-EBOV vaccine study. The animals were originally vaccinated intramuscularly (IM) with a single dose of 1×10^7^ PFU of VSV-EBOV either 28, 21, 14, 7 or 3 days prior to EBOV-Makona strain (10,000 TCID_50_) challenge. The study design is shown in Fig. S2A, and the outcome was previously published^16^. Eleven animals that were protected from EBOV challenge were repurposed for the current vaccine study. Of these, 9 animals were completely protected from disease and never showed EBOV viremia and 2 animals developed mild disease with low level of EBOV viremia^16^. All 11 macaques were rested for ~9 months prior to the VSV-SUDV study start.

### VSV-SUDV protects macaques from SUDV associated clinical disease

Approximately 1 year after the initial VSV-EBOV vaccination, the macaques were divided into two groups. NHPs of one group (n=6) were IM vaccinated with 1×10^7^ PFU of VSV-SUDV and NHPs in the control group (n=5) were IM vaccinated with 1×10^7^ PFU of VSV-MARV. Vaccination with VSV-SUDV and VSV-MARV did not result in any obvious or noticeable adverse effects. After 28 days, all NHPs in both groups were challenged with 10,000 TCID_50_ of SUDV-Gulu by the IM route (Fig. S2A). While none of the VSV-SUDV vaccinated macaques showed any signs of disease in response to challenge and were completely protected, 4 of 5 animals (80%) in the control group developed characteristic clinical signs of SVD and had to be euthanized according to our endpoint scoring criteria (Fig. 1A,B). Control NHPs succumbing to SUDV infection developed lymphocytopenia (Fig. 1C) and high-titer viremia. However, the one surviving control NHP while developing lymphocytopenia was only weakly positive on 6 days post-challenge (DPC) for SUDV RNA but no virus isolation from whole blood was possible (Fig. 1D, E). Compared to the protected NHPs, the 4 control NHPs that succumbed to infection presented with elevated aspartate transaminase (AST), alkaline phosphatase (ALP) and blood urea nitrogen (BUN) (Fig. 1F-H) levels. In addition, these NHPs had hypoalbuminemia (Fig. 1I). All of these clinical chemistry changes are consistent with clinical SVD^17^. The single surviving control NHP showed serum clinical chemistries similar to the protected VSV-SUDV-vaccinated NHPs.

**Figure 1.**
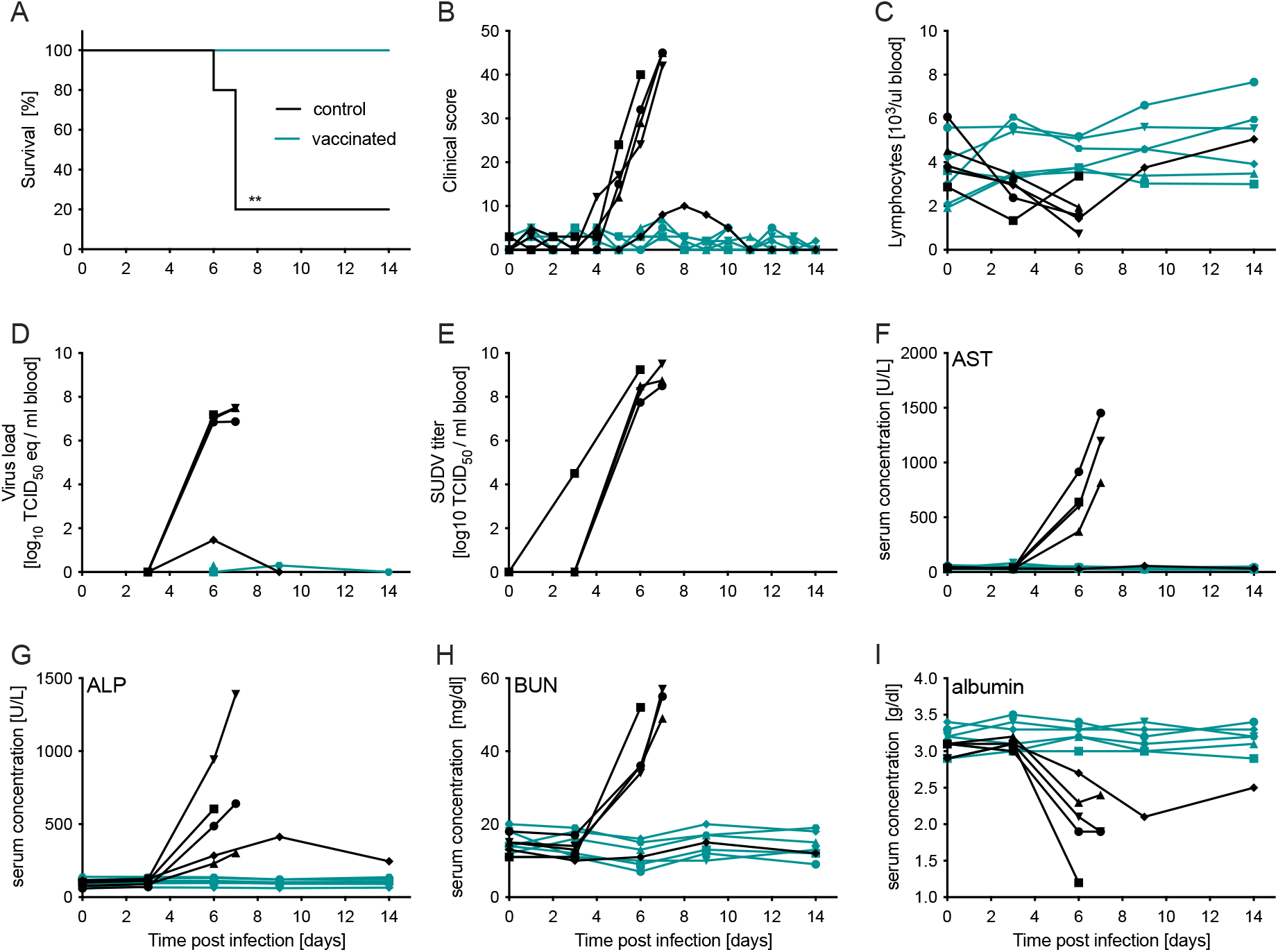
Survival and clinical changes in NHPs after SUDV challenge. Groups of NHPs were vaccinated with VSV-SUDV (n=6) or control vaccine (VSV-MARV; n=5) and challenged 4 weeks later with SUDV. (A) Survival, (B) clinical scores, (C) lymphocytes, viremia by (D) RT-qPCR and (E) titration are shown. Serum levels of (F) aspartate aminotransferase (AST), (G) alanine phosphatase (ALP), (H) blood urea nitrogen (BUN) and (I) albumin were determined. Significant differences in the survival curves were determined performing Log-Rank analysis. Statistical significance is indicated as ** *p*<0.01.

### VSV-SUDV prevents macaques from developing SVD-associated pathology

At the time of euthanasia, the control macaques presented with liver and spleen pathology as previously described for SUDV infections^17^. Histologically the control NHPs demonstrated liver lesions characteristic for SVD including multifocal to coalescing hepatocellular degeneration and necrosis with acute inflammation and abundant micro-fibrin thrombi (Fig. 2A). In the spleen white pulp necrosis and loss with abundant fibrin effacing the red pulp were observed. Immunohistochemical evaluation demonstrated abundant viral antigen associated with these hepatic and splenic lesions (Fig. 2A). High SUDV titers were found in target tissues such as liver, spleen, adrenal glands, lymphoid tissues, urinary bladder and muscle at the injection site (Fig. 2B). Vaccinated NHPs and the single survivor in the control group were necropsied on 40 DPC. All collected tissue samples were essentially normal with no evidence of immunoreactivity in liver or spleen (Fig. 2A).

**Figure 2.**
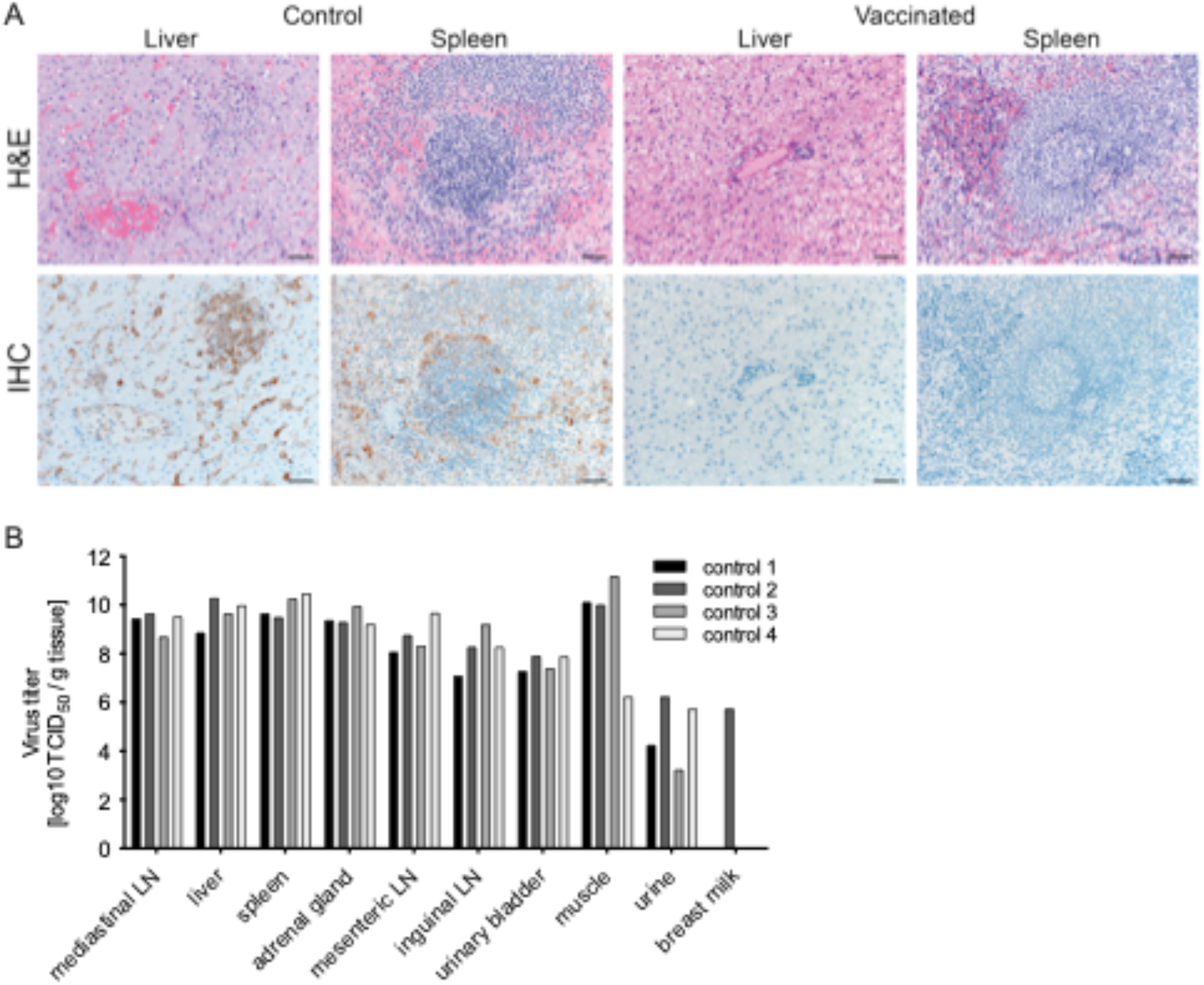
Histopathology after SUDV challenge. (A) At the time of euthanasia liver and spleen were collected (6-8 DPC control; 40 DPC vaccinated), inactivated, processed, and stained with hematoxylin and eosin (H&E). SUDV antigen (VP40) was detected in control NHPs only. Immunoreactive cells are brown. Sections from a representative animal in each group are shown. Magnification 200x; scale bar = 50μm. (B) SUDV titers in control NHPs at the time of euthanasia.

### Pre-existing immunity to EBOV and VSV has limited impact on the development of SUDV GP-specific IgG

Previous studies using the VSV-vectored filovirus vaccines have demonstrated the importance of antigen-specific IgG for protection in NHPs with rather limited contributions from T cell responses^18–20^. Therefore, we focused on the humoral immune response; peripheral blood mononuclear cells (PBMC) for T cell responses were not collected in this study. Prior to VSV-SUDV or VSV-MARV vaccination, all animals still showed an EBOV GP-specific IgG titer of >1:10^3^ (Fig. S2B). The response was significantly boosted with the VSV-SUDV vaccination by more than a magnitude. In contrast, no boosting effect was noticed with the VSV-MARV vaccination (Fig. 3A). The SUDV challenge did not further boost the EBOV GP-specific IgG titers except for the single survivor in the control group which showed a steep booster effect peaking at 14 DPC indicative of an anamnestic response (Fig. 3A).

**Figure 3.**
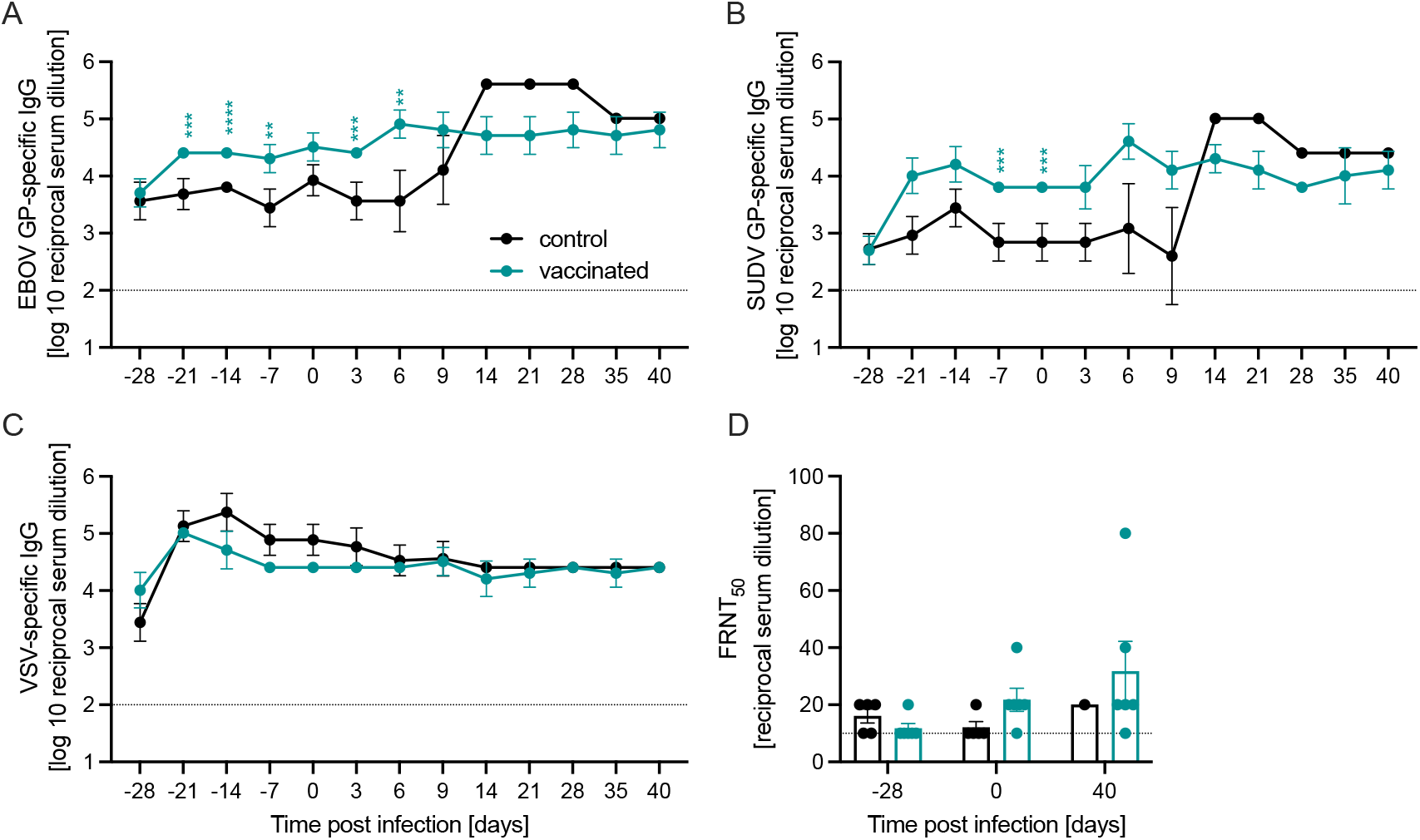
Humoral immune responses after vaccination and SUDV challenge. Serum IgG levels specific for (A) EBOV GP, (B) SUDV GP, and (C) VSV were determined over time. Geometric mean and geometric SD are depicted. Statistical significance was determined by two-way ANOVA with Tukey’s multiple comparisons and is indicated as follows: *p*<0.0001 (****), *p*<0.001 (***), and *p*<0.01 (**). (D) Serum neutralization presented as 50% fluorescence reduction (FRNT50) of GFP-positive cells at the time of vaccination (day-28), challenge (day 0) and euthanasia (day 40; study end). Dotted line presents the limit of detection.

Investigation of the SUDV GP-specific IgG response revealed that all macaques showed limited cross-reactive antibodies from the previous VSV-EBOV vaccination and EBOV challenge prior to VSV-SUDV and VSV-MARV vaccination (Fig. 3B; Fig. S2B). Following VSV-SUDV vaccination, the SUDV GP-specific IgG peaked 14 days after vaccination (titer of >1:10^4^) and was slightly boosted by the SUDV infection reaching its highest level at 6 DPC (1:25,600-1:102,400) (Fig. 3B). The sole survivor among the control animals again showed a steep booster effect following SUDV challenge peaking at 14 DPC indicative of an anamnestic response (Fig. 3B).

When we compared the VSV-specific IgG over the course of this experiment, we again observed peak titers 1-2 weeks after vaccination without a significant difference between the VSV-SUDV study group and VSV-MARV control group (Fig. 3C). These titers stabilized by 0 DPC and remained constant throughout the study.

Analysis of neutralizing immune responses using a VSV-SUDV-GFP-based assay revealed limited neutralizing activity in all vaccinated NHPs at the time of challenge and at study end (Fig. 3D). Remarkably, the serum of the single surviving control NHP demonstrated only a small increase in neutralizing activity on 40 DPC despite the strong increase of SUDV GP-specific IgG responses after challenge.

## Discussion

The current SVD outbreak in Uganda has again shown that the public health response to emerging/reemerging infectious diseases is vulnerable, lacking accredited and stockpiled vaccine and treatment options. Despite great effort and success for EBOV, there still is a general lack of available countermeasures against infections with other human-pathogenic filoviruses. Disappointingly, more than 8 years after the EBOV epidemic in West Africa the situation with vaccines and treatment options against ebolaviruses in general seems similar to when EBOV struck West Africa. Multiple SUDV vaccine candidates (Table S2) and some treatment options have finished preclinical evaluation and are awaiting clinical trials, yet nothing was immediately ready for deployment when this SVD outbreak was declared in September 2022. Clinically, SVD is indistinguishable from EVD and Marburg virus disease, but the pathogens are genetically and antigenically distinct enough to render specific vaccines and monoclonal antibody treatments ineffective against a heterologous infection. Hence, EBOV-specific vaccines, such as VSV-EBOV (Ervebo, Merck), are not expected to be effective against SUDV infections due to lack of cross-protective immune responses as demonstrated in this study. Therefore, we generated a VSV-SUDV vector by simply exchanging the EBOV GP with the SUDV GP to be used as a single-dose, live-attenuated vaccine against SVD. We demonstrated that VSV-SUDV completely protected cynomolgus macaques against lethal challenge with SUDV. Vaccinated animals did not develop viremia, organ tissue viral loads or damage, or clinical disease in strong contrast to the VSV-MARV-vaccinated control NHPs. Interestingly, one control vaccinated NHP survived with a clinical progression similar to the vaccinated NHPs. This may be explained by the published observation that the cynomolgus macaque model for SVD is not uniformly lethal^21^. However, the almost complete lack of SVD parameters in this surviving animal is surprising and interesting and may indicate certain cross-protective immune responses in this animal which is supported by a strong anamnestic IgG response to SUDV challenge. Future studies challenging VSV-EBOV-vaccinated NHPs with SUDV need to confirm or disprove this finding.

This unique study design allowed us to decipher potential cross-reactive or cross-protective immune responses between EBOV and SUDV. All macaques had detectable EBOV GP-specific IgG antibody titers approximately one year after VSV-EBOV vaccination and EBOV challenge (Fig. S2B). They also maintained similar levels of VSV-specific immune responses prior to start of the VSV-SUDV vaccine study. Neither EBOV nor VSV pre-existing immunity hampered the development of SUDV GP-specific humoral immune responses again highlighting the reusability of VSV-ΔG vectors with heterologous glycoproteins as previously shown^22^. In contrast, the VSV-SUDV vaccination boosted the EBOV GP-specific responses likely adding a durability benefit of these protective responses. The SUDV GP-specific immune responses were protective against SUDV challenge whereas the EBOV GP-specific immune responses were not as demonstrated by the control group. Therefore, cross-reactive antibodies are generated between EBOV and SUDV but those are unlikely to cross-protect against heterologous challenge. Although not investigated in this study, it appears that the induced T cell responses to VSV vaccination and EBOV or SUDV challenge cannot overcome the lack of cross-protective antibodies. This may again indicate a minor role of T cell responses to the overall efficacy of VSV-ΔG vectored vaccines^19^.

The study has limitations that need to be addressed by future work. A VSV-EBOV vaccinated group would have been an important control to address the role of cross-protection. However, the study was initially performed to investigate the protective efficacy of VSV-SUDV against SUDV challenge and VSV-MARV vaccination was chosen as control vaccine to limit cross-protective properties. The lack of T cell analysis is a limitation and even though immune response analyses of VSV-vectored filovirus vaccines indicate a minor role of T cells^19^, a complete evaluation of immune responses should include T cell immune responses. Furthermore, the current study does not address the fast-acting efficacy of VSV-SUDV which would have been beneficial for the potential use of this vaccine in ring vaccination strategies for outbreak response. In conclusion, VSV-SUDV completely protects macaques against lethal SUDV challenge and likely acts similarly to other VSV-ΔG-based filovirus vaccines including VSV-EBOV (Ervebo, Merck) and VSV-MARV. The vaccine is a potent candidate for clinical trials including ring vaccination for which this vaccine was developed. Similar to VSV-EBOV the replicating nature of VSV-SUDV likely confers a rapid onset of protective immunity. The lack of cross-protection from VSV-EBOV vaccination and EBOV challenge confirms the ineffectiveness of the licensed VSV-EBOV (Ervebo, Merck) in the current SUDV outbreak. However, the boost of the EBOV GP-specific response through VSV-SUDV vaccination may add a benefit to the durability of protection against EVD which needs to be further assessed. Finally, VSV-SUDV is intended as a vaccine in individuals, such as frontline workers, regardless of previous VSV-EBOV vaccination status.

## Materials & Methods

### Ethics statement

All work involving EBOV and SUDV was performed in the maximum containment laboratory (MCL) at the Rocky Mountain Laboratories (RML), Division of Intramural Research, National Institute of Allergy and Infectious Diseases, National Institutes of Health. RML is an AAALACi-accredited institution. All procedures followed RML Institutional Biosafety Committee (IBC)-approved standard operating procedures (SOPs). Animal work was performed in strict accordance with the recommendations described in the Guide for the Care and Use of Laboratory Animals of the National Institute of Health, the Office of Animal Welfare and the Animal Welfare Act, United States Department of Agriculture. This study was approved by the RML Animal Care and Use Committee (ACUC), and all procedures were conducted on anesthetized animals by trained personnel under the supervision of board-certified clinical veterinarians. The NHPs were observed at least twice daily for clinical signs of disease according to a RML ACUC-approved scoring sheet and humanely euthanized when they reached endpoint criteria.

NHPs were housed in adjoining individual primate cages that enabled social interactions, under controlled conditions of humidity, temperature, and light (12 hours light - dark cycles). Food and water were available *ad libitum*. NHPs were monitored and fed commercial monkey chow, treats, and fruit at least twice a day by trained personnel. Environmental enrichment consisted of commercial toys, music, video, and social interaction. All efforts were made to ameliorate animal welfare and minimize animal suffering in accordance with the Weatherall report on the use of NHPs in research (https://royalsociety.org/policy/publications/2006/weatherall-report/).

### Vaccine vectors

Previously described VSV-based vaccine vectors expressing the EBOV-Kikwit GP (VSV-EBOV)^16^ and VSV-MARV ^20,23^ were used in this study. The VSV-SUDV was generated by cloning the SUDV-Gulu GP gene (GenBank NC_006432.1; 8A version) into the VSV backbone (Fig. S1A top) as previously described for other filovirus GPs ^15^. Similarly, VSV-SUDV-GFP was generated by adding the GFP gene as an additional ORF between the SUDV GP and VSV-L genes (Fig. S1A bottom). Antigen expression was verified by Western blot analysis using anti-EBOV GP (ZGP 42/3.7, 1:10,000; a kind gilt from Ayato Takada, Hokkaido University, Sapporo, Japan), and anti-VSV M (23H12, 1:1,000; Kerafast Inc.).

### Challenge virus

EBOV-Makona Guinea C07 was used as challenge virus in the first study^16^. SUDV-Gulu (GenBank NC_006432.1) was obtained from United States Army Medical Research Institute of Infectious Diseases. The virus was propagated on Vero E6 cells (mycoplasma negative), titered by median tissue culture infectious dose (TDCID50) assay on Vero E6 cells and stored in liquid nitrogen. Deep sequencing revealed no contaminants however, 4 base pair changes (3 of them coding) were noted from this viral passage compared to its reference sequence (Table S3). A target dose of 10,000 TCID_50_ (backtitered as 5,623 TCID_50_) was used for the IM SUDV challenge.

### NHP study design

Eleven male or female cynomolgus macaques, 6-11 years of age and 3.8-9.0 kg at the time of VSV-SUDV vaccination, were used in this study. After the EBOV challenge of the study, NHPs were rested for ~ 9 months and IM-vaccinated with 1× 10^7^ PFU VSV-SDUV or VSV-MARV (control)(Fig. S2A). The NHPs were divided into 2 study groups as outlined in Fig. S2B. All animals received a 1ml IM injection for vaccination into 2 sites in the caudal thighs containing either 1×10^7^ PFU VSV-SUDV (n=6) or 1x 10^7^ PFU VSV-MARV (control; n=5). All NHPs were challenged IM on 0 DPC with 1×10^4^ TCID_50_ SUDV-Gulu into 2 sites in the caudal thighs. Physical examinations and blood draws were performed as outlined in Fig. S2A on −28, −21, −14, −7, 0, 3, 6, 9, 14, 21, 28, and 35 DPC and at euthanasia (day 40 for survivors; humane endpoint for non-survivors). Following euthanasia, a necropsy was performed, and samples of key tissues were collected for analysis.

### Hematology and serum chemistries

Blood cell counts were determined from EDTA blood with the IDEXX ProCyte DX analyzer (IDEXX Laboratories, Westbrook, ME). Serum biochemistry (including AST, ALP, albumin, and BUN) was analyzed using the Piccolo Xpress Chemistry Analyzer and Piccolo General Chemistry 13 Panel discs (Abaxis, Union City, CA).

### SUDV RNA and titer

SUDV RNA copy numbers in EDTA blood samples after challenge were determined using a RT-qPCR assay specific to the SUDV GP; sequences were as follows: forward primer CAAAGGGAAGAATCTCCGACC; reverse primer CAGGGGAATTCTTTGGAACC; probe GGCCACCAGGAAGTATTCGGACC. Blood samples were extracted with QIAmp Viral RNA Mini Kit (Qiagen, Hilden, Germany) according to manufacturer specifications. One-step RT-qPCR was performed with QuantiFast Probe RT-PCR+ROX Vial Kit (Qiagen) on the Rotor-Gene Q (Qiagen). RNA from the SUDV stock was extracted the same way and used alongside samples as standards with known TCID_50_ concentrations. SUDV titers in macaque EDTA blood and tissue samples were determined on VeroE6 cells (mycoplasma negative) using a TCID_50_ assay as previously described for EBOV ^24^. Titers were calculated using the Reed-Muench method ^25^.

### Histology and immunohistochemistry

Necropsies and tissue sampling were performed according to IBC-approved SOPs. Collected tissues were fixed, processed and stained as previously described^26^. Specific anti-VP40 immunoreactivity was detected using a cross-reactive anti-EBOV VP40 antibody (a generous gift by Dr. Yoshihiro Kawaoka, University of Wisconsin-Madison) at a 1:1,000 dilution. All tissue slides were evaluated by a board-certified veterinary pathologist.

### Assessment of humoral immune response

Post-challenge NHP sera were inactivated by γ-irradiation (4 MRad), a well-established method with minimal impact on serum antibody binding ^27,28^, and removed from the MCL according to SOPs approved by the RML IBC. Titers for IgG specific to EBOV GP or SUDV GP were determined in endpoint dilution ELISAs using recombinant EBOV GPΔTM (#0501-001; IBT Bioservices, Rockville, MD) or recombinant SUDV GPΔTM (#0502-001; IBT Bioservices) as described previously ^29^. Levels of VSV-specific IgG were determined by endpoint dilution ELISA using concentrated VSV wildtype particles lysed with 0.01% triton-X100 in PBS as antigen.

Neutralization of irradiated and heat-inactivated serum samples were assessed in Vero E6 cells. Briefly, cells were seeded in 96-well round-bottom plates for 24 hours. On the day of neutralization, serial dilutions of heat-inactivated serum samples were performed in DMEM supplemented with 2% FBS, penicillin/streptomycin, and L-glutamine. Each plate contained a negative serum control, cell-only control, and virus-only control. VSV-SUDV-GFP was added to each well of the serum dilution plate and the serum-virus mix was incubated at 37°C for 1-hour. The mix was added to the cells and incubated at 37°C for 24 hours. The cells were then fixed with 4% paraformaldehyde at room temperature for 15-minutes and centrifuged at 600 x g for 5-minutes at room temperature. The supernatant was discarded and FACS+EDTA buffer was added. Samples were run on the FACSymphony A5 Cell Analyzer (BD Biosciences, Mississauga, ON, Canada) and FITC MFI was measured. Data were analyzed using FlowJo V10.

### Statistical analysis

Statistical analysis was performed in Prism 9 (GraphPad). Data presented in Fig. 3 were analyzed by two-way ANOVA with Tukey’s multiple comparison to evaluate statistical significance at all timepoints between all groups. Significant differences in the survival curves shown in Fig. 1A were determined performing Log-Rank analysis. Statistical significance is indicated as *p*<0.0001 (****), *p*<0.001 (***), *p*<0.01 (**), and * *p*<0.05.

## Acknowledgments

We thank members of the Rocky Mountain Veterinary Branch, NIAID for supporting the NHP study. We are grateful to the United States Army Medical Research Institute of Infectious Diseases for sharing their virus isolate.

## Author Contributions

A.M. and H.F. conceived the idea, designed the study, and secured funding. A.M., F.F., and P.W.H. conducted the NHP study. A.M., P.F., and G.S. collected and processed samples, performed assays, and analyzed the data. A.M. and H.F. wrote the manuscript with input from all authors. All authors approved the manuscript.

## Data availability

All data is presented in the paper and available from the corresponding author upon request.

---

**Table S1.**
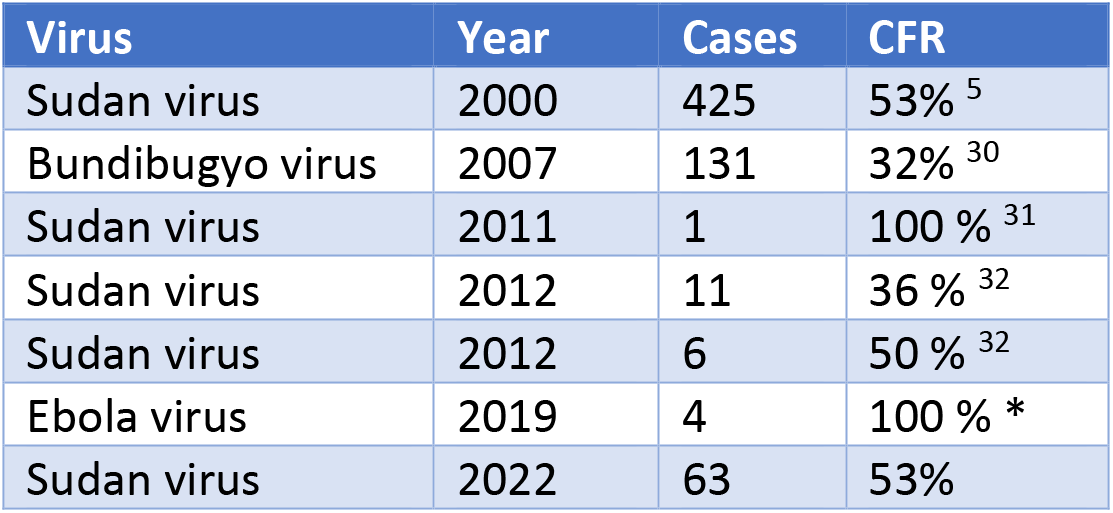
Outbreaks of Ebola disease in Uganda. CFR case fatality rate. *These 4 cases occurred during the 2018-2020 Ebola virus outbreak in The Democratic Republic of the Congo (DRC). They were attributed to cross-border movement from DRC and are accounted for in the DRC outbreak statistics.

| Virus | Year | Cases | CFR |
| --- | --- | --- | --- |
| Sudan virus | 2000 | 425 | 53%<sup>5</sup> |
| Bundibugyo virus | 2007 | 131 | 32%<sup>30</sup> |
| Sudan virus | 2011 | 1 | 100 %<sup>31</sup> |
| Sudan virus | 2012 | 11 | 36 %<sup>32</sup> |
| Sudan virus | 2012 | 6 | 50 %<sup>32</sup> |
| Ebola virus | 2019 | 4 | 100 % * |
| Sudan virus | 2022 | 63 | 53% |

**Table S2.** SUDV vaccines with preclinical efficacy in NHPs. VLPs virus-like particles. GP glycoprotein. VEEV Venezuelan equine encephalitis virus. MVA Modified vaccinia Ankara. TAFV Taï Forest virus. IM intramuscular. D days. *days after vaccinations were completed. IAVI International AIDS Vaccine Initiative.

| Vaccine | Challenge Virus and route | Vaccine doses | Time between doses [d] | Time to Challenge [d]* | Survival [%] | Developer | Ref. |
| --- | --- | --- | --- | --- | --- | --- | --- |
| **Subunit** |  |  |  |  |  |  |  |
| VLPs + adjuvant | SUDV Boniface IM | 2 | 42 | 28 | 100 | USAMRIID | 33 |
| SUDV GP, EBOV GP + adjuvant | SUDV Gulu IM | 3 | 21 | 28 | 100 | U Hawaii | 34 |
| SUDV GP + adjuvant | SUDV Gulu IM | 3 | 21 | 28 | 100 | U Hawaii | 34 |
| **Replicon** |  |  |  |  |  |  |  |
| VEEV-SUDV GP | SUDV Boniface IM | 1 | n/a | 28 | 100 | USAMRIID | 35 |
| VEEV-SUDV GP + VEEV-EBOV GP | SUDV Boniface IM | 1 | n/a | 28 | 100 | USAMRIID | 35 |
| VEEV-SUDV GP | SUDV Boniface aerosol | 1 | n/a | 28 | 0 | USAMRIID | 35 |
| VEEV-SUDV GP | SUDV Boniface aerosol | 2 | 28 | 56 | 100 | USAMRIID | 35 |
| **rec. Adenovirus** |  |  |  |  |  |  |  |
| CAdVax-EBO7 | SUDV Boniface IM | 2 | 120 | 42 | 100 | UASMRIID | 36 |
| CAdVax-EBO7 | SUDV Boniface aerosol | 1 | n/a | 28 | 67 | UASMRIID | 36 |
| CAdVax-EBO7 | SUDV Boniface aerosol | 2 | 71 | 28 | 100 | UASMRIID | 36 |
| Ad26.Filo (EBOV, SUDV, MARV) | SUDV Gulu IM | 2 | 28 | 28 | 75 | Janssen | 37 |
| Ad26.Filo + Ad35.Filo (EBOV, SUDV, MARV) | SUDV Gulu IM | 2 | 28 | 28 | 75 | Janssen | 37 |
| Ad5-SUDV | SUDV Gulu IM | 1 | n/a | 28 | 50 | Janssen | 37 |
| **rec. Adenovirus + MVA** |  |  |  |  |  |  |  |
| Ad26.Filo + MVA-BN-Filo (EBOV, SUDV, MARV, TAFV) | SUDV Gulu IM | 2 | 56 | 28 | 100 | Janssen | 38 |
| MVA-BN-Filo + Ad26.Filo (EBOV, SUDV, MARV, TAFV) | SUDV Gulu IM | 2 | 56 | 28 | 60 | Janssen | 38 |
| Ad26.Filo + Ad35.Filo (EBOV, SUDV, MARV) | SUDV Gulu IM | 2 | 56 | 28 | 100 | Janssen | 38 |
| **rec. Vesicular stomatitis virus** |  |  |  |  |  |  |  |
| VSV-EBOV, VSV-SUDV, VSV-MARV | SUDV Boniface IM | 1 | n/a | 28 or 59 | 100 | IAVI | 10 |
| Trivalent VesiculoVax | SUDV Gulu IM | 1 | n/a | 28 | 100 | Auro Vaccines | 39 |
| Quadrivalent VesiculoVax | SUDV Gulu IM | 1 | n/a | 28 | 100 | Auro Vaccines | 40 |

**Table S3.** Base pair changes SUDV-Gulu RML stock versus GenBank reference. NP nucleoprotein; VP40 virion protein 40; GP glycoprotein; bp base pair.

| Gene | Location (bp) | Stock | Nucleotide | Amino Acid | Coding Seq |
| --- | --- | --- | --- | --- | --- |
| NP | 2591 | NC_006432.1 | T | Y | Yes |
|  |  | RML stock | C | H |  |
|  | 2618 | NC_006432.1 | C | L | Yes |
|  |  | RML stock | T | F |  |
| VP40 | 5320 | NC_006432.1 | C | L | Yes |
|  |  | RML stock | T | K |  |
| GP | 5902 | NC_006432.1 | G | n/a | No |
|  |  | RML stock | T |  |  |

**Figure S1.**
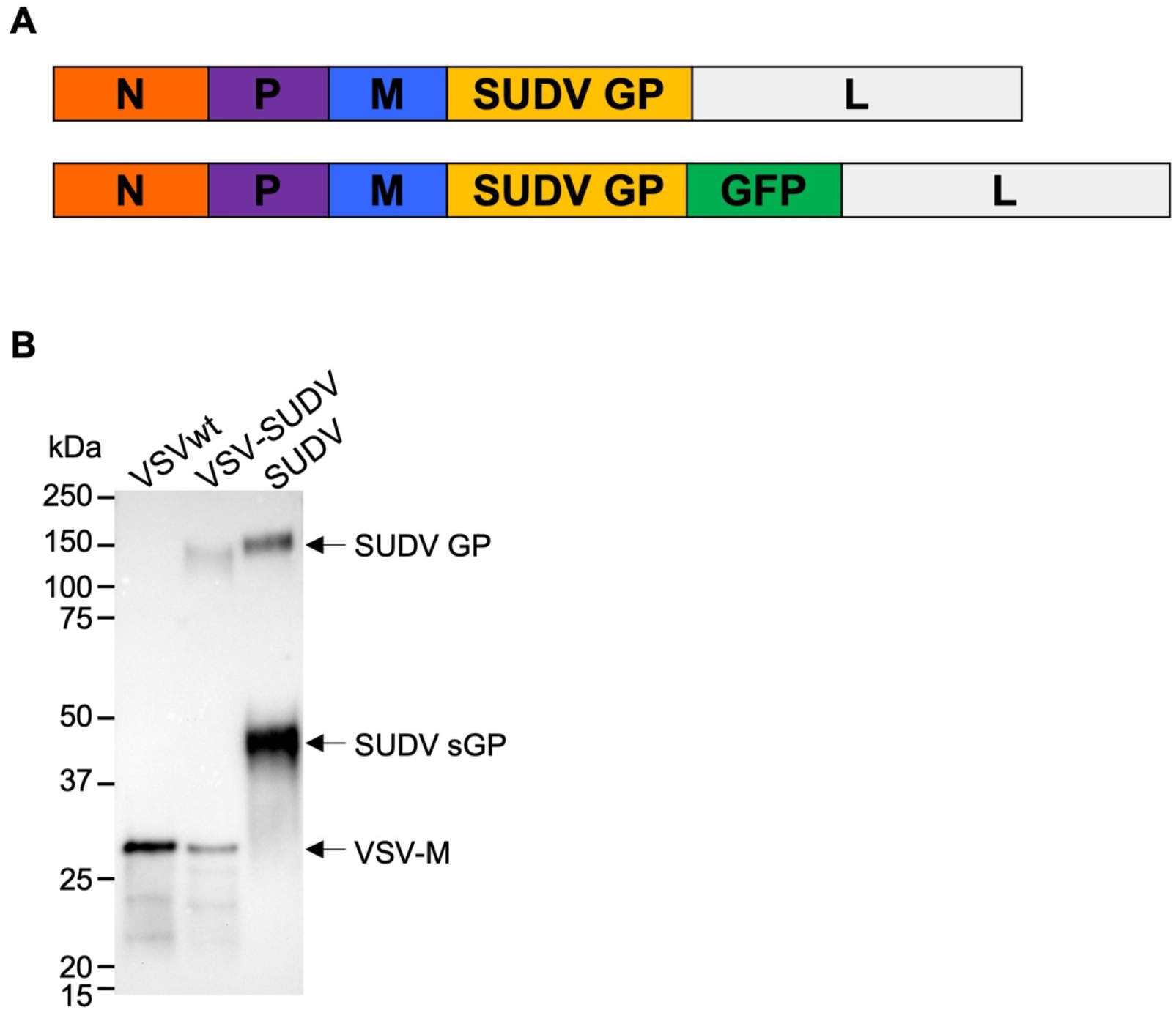
Characterization of VSV-SUDV. (A) Schematic of the placement of the viral antigen in the VSV genome. Top shows the VSV-SUDV used for vaccination. The VSV-SUDV-GFP (bottom) was used for neutralization testing. N nucleoprotein, P phosphoprotein, M matrix protein, G glycoprotein, L polymerase, SUDV GP Sudan virus glycoprotein, GFP green fluorescent protein. (B) Antigen expression by Western blot. SUDVGP (~150 kDa), SUDV soluble GP (sGP; ~45 kDa), and VSV-M (~30 kDa) were detected by monoclonal antibody staining.

**Figure S2.**
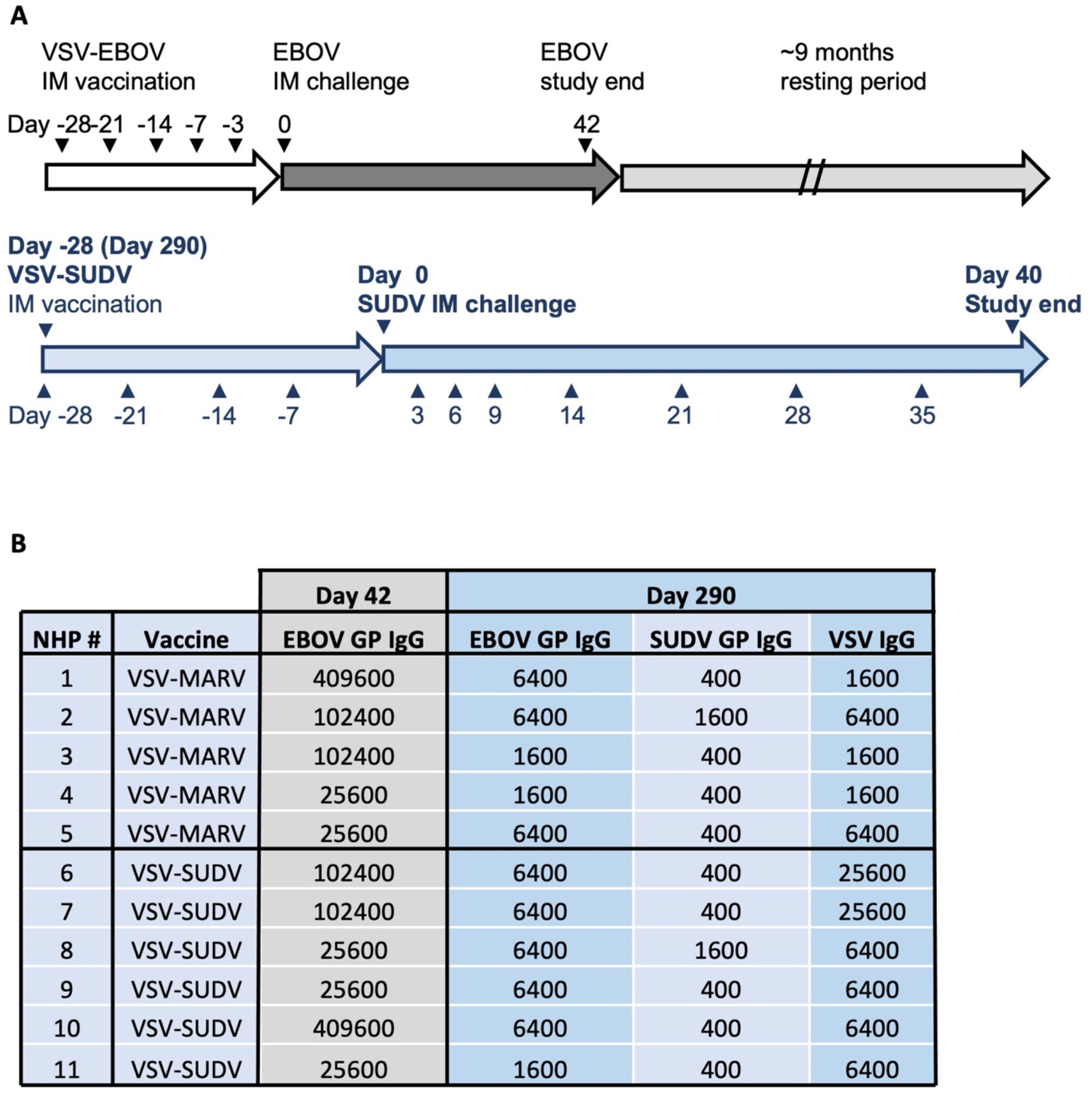
Study outline. (A) Timeline of the EBOV and SUDV parts of this study. (B) Details on NHP vaccination with VSV-EBOV and EBOV GP-specific IgG titers at the end of the EBOV challenge study (day 42). On day 290, a serum sample was collected before the NHPs were vaccinated with VSV-SUDV or VSV-MARV (control). Reciprocal endpoint titers of IgG specific for SUDV GP, EBOV GP, or VSV are listed.

